# A type 2 diabetes disease module with a high collective influence for Cdk2 and PTPLAD1 is localized in endosomes

**DOI:** 10.1101/341693

**Authors:** Martial Boutchueng-Djidjou, Pascal Belleau, Nicolas Bilodeau, Suzanne Fortier, Sylvie Bourassa, Arnaud Droit, Sabine Elowe, Robert L. Faure

## Abstract

Despite the identification of many susceptibility genes our knowledge of the underlying mechanisms responsible for complex disease remains limited. Here, we identified a type 2 diabetes disease module in endosomes, and validate it for functional relevance on selected nodes. Using hepatic Golgi/endosomes fractions, we established a proteome of insulin receptor-containing endosomes that allowed the study of physical protein interaction networks on a type 2 diabetes background. The resulting collated network is formed by 313 nodes and 1147 edges with a topology organized around a few major hubs with Cdk2 displaying the highest collective influence. Overall, 88% of the nodes are associated with the type 2 diabetes genetic risk, including 101 new candidates. The Type 2 diabetes module is enriched with cytoskeleton and luminal acidification –dependent processes that are shared with secretion-related mechanisms. We identified new signaling pathways driven by Cdk2 and PTPLAD1 whose expression regulate the association of the insulin receptor with TUBA, TUBB, the actin component ACTB and the endosomal sorting markers Rab5c and Rab11a. Therefore, the interactome of internalized insulin receptors reveals the presence of a type 2 diabetes disease module enriched in new layers of feedback loops required for insulin signaling, clearance and islet biology.

**Author Summary:** According to the local hypothesis each complex disease can be linked to a well-defined network called the disease module. A disease module can be defined by the topological properties of protein interaction networks. Given the complexity of the whole interaction map the existence of such disease modules remains largely to be tested. Here, we found a type 2 diabetes disease module in insulin receptor-containing endosomes. The disease module contains new pathways that are associated with both insulin signaling, clearance and secretion. Its co-functionality with islets biology may provide a mechanistic rationale for the exploration of personalized medicine and elaborate new drugs.

## Introduction

The insulin receptor (IR) belongs to the receptor tyrosine-kinase (RTK) family of cell-surface receptors [1, 2]. Early work on insulin and epidermal growth factor (EGF) revealed the presence of signaling molecules in hepatic endosomes fractions [3]. The concept of endosomal signaling is now well established [4-6], but the rules underlying IR trafficking and signaling compared with those underlying the EGF receptor (EGFR) remain relatively unknown; this may be because proper insulin signaling and trafficking correlate with the maintenance of cell polarity [7].

Type 2 diabetes (T2D) is the result of a chronic energy surplus [8] coupled with a strong hereditary component. Estimates for the heritability of T2D range from 20 to 80% with a sibling relative risk of approximately 2, with obesity being an important driver in every population. The detailed genetic architecture of T2D was recently elucidated, and unlike type 1 diabetes (T1D) where the genetic risk is mostly concentrated in the HLA region, the genetic component explaining part of the heritability of T2D is primarily due to a combination of numerous common variants of small effect scattered across the genome [9- 11]. T2D is characterized by both resistance to the action of insulin and defects in insulin secretion; the former has been an important motivating factor in the exploration of insulin signaling [1, 2]. Previous efforts to demonstrate that the genes mapping close to T2D risk loci are enriched for established insulin signaling pathways, however met with limited success; the most robust finding to date implicates seemingly unrelated cellular mechanisms, the majority of which affect insulin secretion and beta cell function [10-14].

An accumulation of proteins associated with T2D was previously observed in the interactome of the IRs endocytosed in an hepatic Golgi/endosomes fraction [15], suggesting the existence of a disease module at this intracellular locus that could help to further understand IR routing mechanisms, the primary mechanisms of the disease and drive the development of rational approaches for new therapies [16, 17]. Here, starting from a proteome of IR-containing endosomes to narrow the space search and the construction of a T2D-protomodule using validated genes, we reveal the presence of a T2D disease module with functional relevance both to insulin targets and insulin producing cells.

## Results

### Recycling Rabs, V-ATPase subunits, tyrosine phosphatases, and cell cycle proteins shape IR-containing endosomes

To determine the proteomic environment of the internalized IRs, we performed a survey of IR-containing endosomes fractions. We started with a mixed Golgi/endosomes fraction (G/E) using a single dose of insulin (1.5 µg/100 g body weight [b.w.]) that resulted in 50% saturation of rat liver receptors. Fractions were prepared at the 2-minute time peak of IR accumulation and the 15-minute 50% decline time [15, 18] to collect a larger proteome. Freshly prepared fractions were then incubated with anti-IR (β-subunit)-coated magnetic beads, and endosomes were collected with a magnet [19, 20]. We identified a total of 620 proteins with high confidence (named IREP: IR Endosome Proteome, Fig 1A and S1Table). Gene ontology (GO) analysis revealed enrichment of proteins involved in trafficking and signaling (MGI database; Biological Network Gene Ontology (BINGO) tool). These were primarily represented by coat-forming elements, small GTPases, components of the actin cytoskeleton, microtubules and motor proteins of the microtubule cytoskeleton and regulators of the cell cycle (Fig 1A, B left panel). Immunoblotting analysis confirmed the peak of IR accumulation occurring at 2 minutes post-insulin injection (Fig 1B right panel). The protein PTPLAD1 (*HACD3*), previously observed to be associated with the IR in G/E fractions after insulin stimulation [15], was also detected here at 15 minutes post-insulin injection (Figure 1B, right panel). Consistent with the presence of sets of Rabs [19, 21] and the highly recyclable fate of IRs in the liver compared with the low-recycling receptor EGFR in liver [3], thirteen Rabs were identified, all involved in transport from early to recycling endosomes (Rab22a, 2 minutes post-insulin injection) or late recycling endosomes (Rab11a, Rab17) [22]. Rab8a, reported to act exclusively in the trans-Golgi network to plasma membrane transport, was identified at 15 minutes post-insulin injection (Fig 1A). Other Rabs identified at both times (Rab6a, Rab5c, Rab1a, Rab2b, Rab11b, Rab14, Rab1b, Rab7a and Rap1b) are all implicated in recycling, transcytotic or Golgi transport events [19, 22]. Among signaling proteins, the transmembrane protein tyrosine phosphatase (PTP) of the R subfamily [23], PTPRF (also named leukocytes antigen-related, LAR) was identified (Fig 1A). PTPRs are generally associated with IR tyrosine dephosphorylation [24-27], acting preferentially on the juxtamembrane sites Y960 and Y1146 located in the IR activation loop [25, 27]. The putative PTP Dnajc6 (also called auxillin) is a chaperone involved in clathrin-mediated endocytosis of EGFR [28, 29]. PTPN6 (SHP-1) is a known IR regulator in the liver [30]. The large representation of regulators of the cell cycle was less expected but is consistent with the attenuation of endocytosis during cell division [31]. The proton translocation machinery necessary to achieve optimal lumenal acidic pH is also particularly well represented by the identification of V-ATPase subunits (ATPv1a, ATPv1b2, ATPv1f, Atpv1e1, ATPv0a1; 2 minutes post-insulin injection) (Fig 1A and B left panel, S1Table). Efficient acidification by V-ATPase is particularly important for the ligand dissociation-degradation sequence according to the law of mass action and is specific to insulin in contrast with EGF or prolactin complexes. This sequence is followed by a rapid recycling of the freed IR under the concerted action of endosomal protein tyrosine phosphatases (PTPs), thus supporting efficient circulating clearance [3, 32].

**Figure 1.**
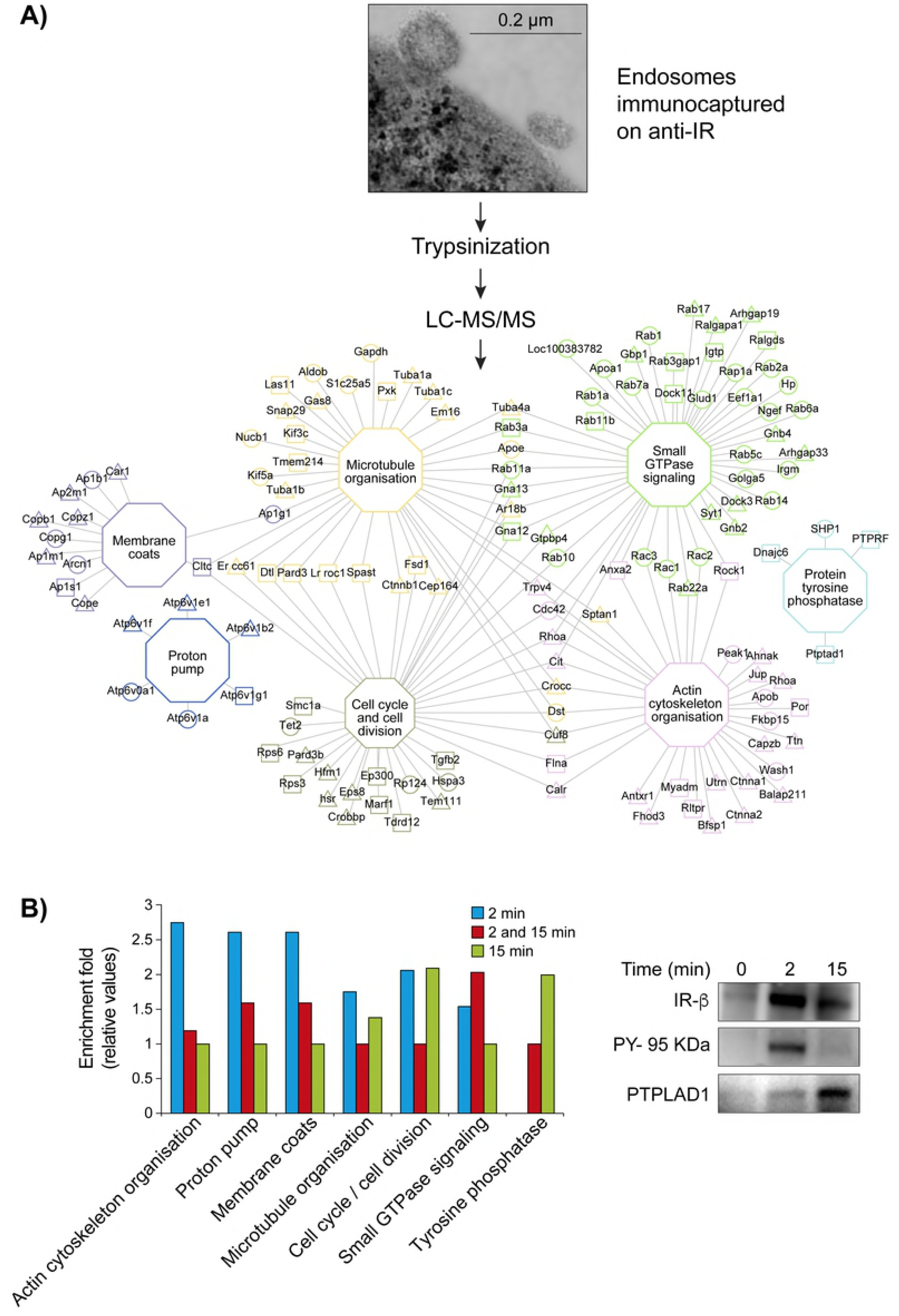
Network of enriched cellular processes in IR-containing endosomes. (A) Workflow of network construction: Inbound endosomal proteins (IREP) were classified into major functional groups according to the MGI database and using the tool BINGO. The triangles (2 minutes) and the squares (15 minutes) are indicative of the insulin post-injection time before endosomal preparation. The circles indicate proteins identified at both times. The hexagonal nodes and their respective border paints represent the functional groups associated linked proteins. Proteins associated with more than one functional group have the border paints of the most statistically significant functional group (Table S 1). (B) (left panel), Comparative enrichment profiles of trafficking proteins according to the insulin post-injection time. (right panel), The bound fraction (100 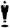 mg of protein) was blotted and pieces were incubated with antibodies against IR (95 kDA 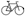-subunit), phosphotyrosine (PY-20, PY-95 kDA) and PTPLAD1.

### Genes at risk for type-2 diabetes form a proto-module enriched for transport and oxygen species regulation

Most of the established T2D genes are supported by low and high probability GWAS signals of their identified variants [10, 11]. To verify if the IREP is associated with T2D, we used complementary data sources (DIAGRAM consortium, SNPs provided in replicated GWAS from the NHGRI-EBI GWAS catalog and GWAS Central portal, source S2Table) to compile a list of 452 T2D and associated trait genes on the basis of single-nucleotide polymorphisms (SNPs) identified in their genomic loci (diabetes-associated gene: DAG; p-value < 5 × 10^−8^; S3Table). This list also contains relevant genes associated with T2D Mendelian traits described in the OMIM database and tagged with the symbol (3) indicative of known molecular associations (S3Table, sheet OMIM). To reduce false-positive associations, the 452 DAG products were validated in a physical protein interaction network (PPIN) [16, 33]. We gathered physical protein-protein interaction data from the Biological General Repository Interaction Datasets (BIOGRID), the human interactomes I and II generated with Y2H systems from the Center for System Biology (CCSB) interactome, Intact, Reactome, Database of Interacting Proteins (DIP, UCLA), HitPredict databases or from the Human Proteins Repository Database (HPRD). The network was visualized with Cytoscape [34]. The 452 DAG products formed a PPIN of 184 proteins and 309 interactions we called the *proto-T2D module* (Fig 2A and S4Table, sheet T2DN-protomodule). The proto-T2D module is made up essentially of protein coding-genes from OMIM (26%), GWAS variants with a p-value < 1×10^−8^; 69%, and GWAS variants with 5 × 10^−8^ < p-value <1 × 10^−8^; 5%, (Fig 2B). It displays a scale-free topology relying on a few hubs of large size such as HNF4A surrounded by a majority of the peripheral nodes (more than 38% of the nodes have only one interactor) [16, 17] (Fig 2A and C; S5Table, sheet-ProtoT2Dmodule). Protein transport (p < 1.82 × 10^−34^) and response to oxygen-containing compounds (p < 1.32 × 10^−32^) are the most enriched cellular processes identified by a gene ontology (GO) analysis with the presence of trafficking proteins (Rab5b, RABEPP1, RABEPP2) and transcription factors from the HNF (HNF4A, HNF1A or HNF1B), FOX (FOXO3, FOXA2) and TCF families (TCF7L2, TCF4, and TCF19). Signaling modules associated with insulin sensitivity are also present (INSR, IRSs, GRB14, PTPN1) (Fig 2A; S5Table, sheet GO analysis-T2D-protomodule). The affinity for these biological processes is supported by the enriched subcellular component analysis which revealed an accumulation of the coding genes associated with risk for T2D in endosomes and endoplasmic reticulum among the major genes. Proteins from the histocompatibility complex are also among the most significant clusters in the T2D-protomodule (Fig 2A; S5Table, sheet GO analysis-T2D-protomodule).

**Figure 2.**
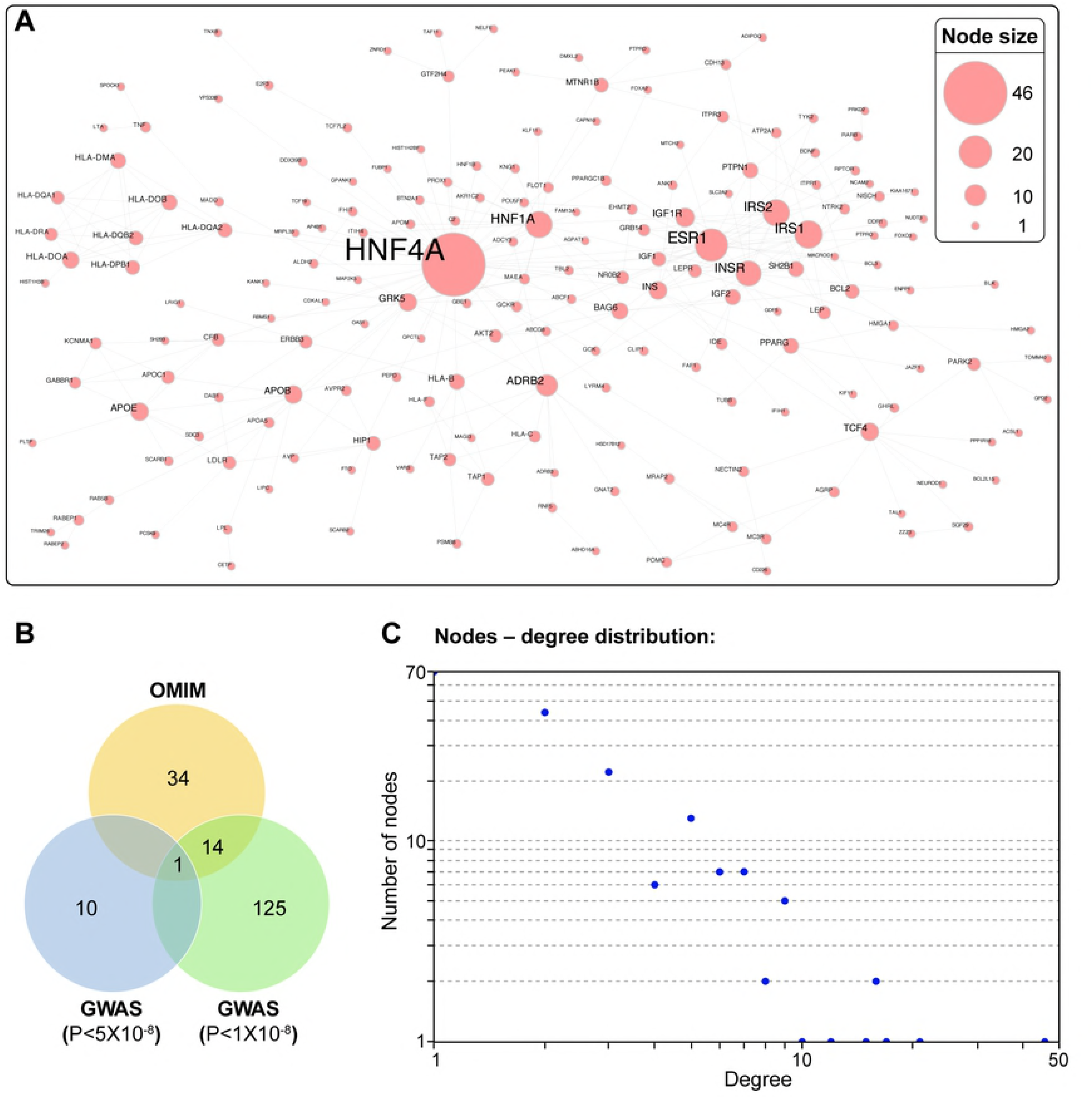
Diabetes-associated genes form a protomodule. (A) Overall, 452 diabetes-associated gene (DAGs; GWAS p value < 5 × 10^−8^ and OMIM; Table S3) products form a PPIN of 184 proteins and 309 interactions (Tables S4, S5) termed T2D-protomodule. (B) In total, 10 % (11/102) of the high-confidence DAGs with a probability less than 5 × 10^−8^, 53 % of the DAGs with a probability less than 1 × 10^−8^ (141/266) and 49 % of the OMIM genes (49/84) are recovered in the proto-T2D module, showing a tendency to select the highest level of reliability. (C) Nodes – degree distribution: More than 38% of nodes (70 nodes) in the proto-T2D module are peripheral with a minority of hubs from transcription factor families. The general topology of the protomodule is characteristic of a disease network with the presence of few central hubs of large size, surrounded by numerous peripheral hubs of smaller size (Table S5).

Overall, 62% of the 452 selected DAGs are disconnected (267/452). Five factors likely contribute to this fragmentation as follows:

i. True lack of binary or indirect physical interaction.
ii. Interactome incompleteness [33].
iii. False positives (not all genes have a known mechanistic association with the disease) and genes associated with late complications of the disease.
iv. The T1D-T2D paradox and disease classification [9].
v. Missing heritability [10, 11].

### A total of 101 high confidence candidate genes for type 2 diabetes risk are identified in IREP

Genes that fall within one of the known disease loci and whose protein products interact with a known risk factor are predicted to be 10-fold enriched in a true disease gene. By considering the cellular localization as well, the network information leads to a 1000-fold enrichment over random genes [16, 35]. We used a combination of approaches to identify candidate genes confidently associated with diabetes traits. The candidates were grouped and ranked according to i) their topological proximity with the 184 previously validated DAG “seeds” in the PPIN approach, ii) the probability of colocalization in the same subcellular locus (S6Table), iii) the identification of proximal variants correlating with the diabetes GWAS signal by fine-mapping analysis (S7Table) and iv) the similarity of gene expression regulation with the 184 seeds of the proto-T2D-module (S8Table), see Methods. A total of 246 nonredundant IREP coding genes are associated with diabetes traits when considering each of the approaches individually. Of these, 38 were validated by at least three of the approaches mentioned previously. This list includes the *Cdk2* gene which is located in the risk area composed of 4 blocks in strong LD around the T2D SNP rs2069408 (Fig S1A; S7Table). ATIC, which was previously observed to be associated with the IR in endosomes together with PTPLAD1 and AMPK [15]; PTPN6 (SHP-1); the small GTPases of Rab the family (Rab14, and Rab5c); and a series of coat components (i.e., AP complexes, CAV1, COPA, SEC23A and SEC24C) are also present (Table 1). Sixty-three other IREP coding genes are shown to have reliable association with diabetes after validation with any two approaches. This list includes the *HACD3* (PTPLAD1) gene, located in a risk area (Fig S1B; S7Table), that was also previously associated with T2D in human islets [36]. PRKAA1 (AMPK); MTHFD1; the Cdk2 regulators (CDKN1B, CCNE1); a V-ATPase subunit (ATP6VA1); the small GTPase Rab1a, Rab1b and Rab8a; several coat components; the putative tyrosine phosphatase DNAJC6; ACTB and TUBA also fall into this category (S9Table, candidates). IREP is also enriched in genes associated with the T2D risk with 15 of the 184 validated DAGs from the human genome being identified indicating a nonrandom concentration of diabetes genes variants during IR endocytosis (p-value of 3.44 × 10^−4^; hypergeometric test) (S4Table, sheet IREP-HUGO).

**Table-1.** List of candidates. Thirty-eight IREP coding genes are validated for association with diabetes traits by at least three out of four distinct approaches. (PPIN) protein-protein interactions network. (GO) Gene Ontology, Subcellular co-localization. (GWAS) fine-mapping. (COEXPRESSION) same expression pattern.

| CANDIDATES/<br>SUBSTRATES | CANDIDATES NAMES | UPSTREAM KINASES IN IREP | VALIDATION |
| --- | --- | --- | --- |
| CDK2 | Cyclin-Dependent Kinase 2 | CDK2 | PPIN/GO/GWAS/COEXPRESSION |
| B2M | Beta-2-Microglobulin | - | PPIN/GO/COEXPRESSION |
| ATP2A2 | Sarcoplasmic/Endoplasmic Reticulum Calcium Atpase 2 | AMPKA1/CAMKK2/CDK2/ROCK1 | PPIN/GO/COEXPRESSION |
| CTNNB1 | Catenin Beta-1 | AMPKA2/CAMKK2/CDK2/CIT/INSR/ROCK1 | PPIN/GO/GWAS |
| GNB4 | Guanine Nucleotide-Binding Protein Subunit Beta-4 | CAMKK2/ROCK1 | PPIN/GO/GWAS |
| HSPA8 | Heat Shock Protein Family A (Hsp70) Member 8 | AMPKA1/CAMKK2/CDK2/CIT/INSR | PPIN/GO/GWAS |
| RAB14 | Ras-Related Protein Rab-14 | AMPKA1 | PPIN/GO/GWAS |
| SEC24A | Protein Transport Protein Sec24a | AMPKA1/AMPKA2/CAMKK2/CDK2 | PPIN/GO/GWAS |
| SEC31A | Protein Transport Protein Sec31a | AMPKA1/AMPKA2/CDK2/CIT/INSR | PPIN/GO/GWAS |
| TFRC | Transferrin Receptor Protein 1 | CAMKK2/CDK2/CIT/INSR | PPIN/GO/GWAS |
| ALB | Albumin | - | PPIN/GO/COEXPRESSION |
| AP1B1 | Ap-1 Complex Subunit Beta-1 | CAMKK2/CDK2/CIT | PPIN/GO/COEXPRESSION |
| AP1G1 | Ap-1 Complex Subunit Gamma-1 | AMPKA2/CAMKK2/CIT | PPIN/GO/COEXPRESSION |
| AP1M1 | Ap-1 Complex Subunit Mu-1 | AMPKA1/CDK2/CIT/INSR/ROCK1 | PPIN/GO/COEXPRESSION |
| AP1S1 | Ap-1 Complex Subunit Sigma-1a | CDK2 | PPIN/GO/COEXPRESSION |
| APOC2 | Apolipoprotein C-II | - | PPIN/GO/COEXPRESSION |
| ATIC | Bifunctional Purine Biosynthesis Protein Purh | CAMKK2/INSR/ROCK1 | PPIN/GO/COEXPRESSION |
| AP2M1 | Ap-2 Complex Subunit Mu | AMPKA1/CAMKK2/CDK2/CIT | PPIN/GO/COEXPRESSION |
| CALR | Calreticulin | AMPKA2/CDK2/ERBB4 | PPIN/GO/COEXPRESSION |
| CAV1 | Caveolin-1 | INSR | PPIN/GO/COEXPRESSION |
| CD74 | Hla Class II Histocompatibility Antigen Gamma Chain | AMPKA1/CAMKK2/INSR/ROCK1 | PPIN/GO/COEXPRESSION |
| CLTC | Clathrin Heavy Chain 1 | AMPKA2/CAMKK2/CDK2/CIT/INSR/ROCK1 | PPIN/GO/COEXPRESSION |
| EEF1A1 | Elongation Factor 1-Alpha 1 | CDK2 | PPIN/GO/COEXPRESSION |
| FGA | Fibrinogen Alpha Chain | AMPKA2/CAMKK2/CDK2/CIT/INSR | PPIN/GO/COEXPRESSION |
| GNAI2 | Guanine Nucleotide-Binding Protein G | CAMKK2/CIT/INSR | PPIN/GO/COEXPRESSION |
| HPX | Hemopexin | CDK2/CIT/ERBB4/INSR | PPIN/GO/COEXPRESSION |
| JUP | Junction Plakoglobin | AMPKA2/CaMKK2/CDK2/ERBB4 | PPIN/GO/COEXPRESSION |
| LRP1 | Low-Density Lipoprotein Receptor-Related Protein 1 | AMPKA1/AMPKA2/CaMKK2/CDK2/CIT/ERBB4/InsR/ROCK1 | PPIN/GO/COEXPRESSION |
| PTPRF | Receptor-Type Tyrosine-Protein Phosphatase F | AMPKA1/AMPKA2/CAMKK2/CDK2/ERBB4/INSR | PPIN/GO/COEXPRESSION |
| RAB5C | Ras-Related Protein Rab-5c | CAMKK2/CDK2/CIT/INSR | PPIN/GO/COEXPRESSION |
| RAP1A | Ras-Related Protein Rap-1a | CDK2/ERBB4 | PPIN/GO/COEXPRESSION |
| SDC1 | Syndecan-1 | AMPKA1/CDK2/INSR | PPIN/GO/COEXPRESSION |
| SEC23A | Protein Transport Protein Sec23a | AMPKA1/CDK2/INSR | PPIN/GO/COEXPRESSION |
| SEC24C | Protein Transport Protein Sec24c | AMPKA2/CAMKK2/CDK2/CIT/INSR | PPIN/GO/COEXPRESSION |
| ATP5B | Atp Synthase Subunit Beta, Mitochondrial | AMPKA1/AMPKA2/CDK2/CIT | PPIN/GWAS/COEXPRESSION |
| COPA | Coatomer Subunit Alpha | AMPKA1/AMPKA2/CDK2/ERBB4 | PPIN/GWAS/COEXPRESSION |
| GBF1 | Golgi-Specific Brefeldin A-Resistance Guanine Nucleotide Exchange Factor 1 | AMPKA1/AMPKA2/CAMKK2/CDK2/ERBB4/ROCK1 | PPIN/GWAS/COEXPRESSION |
| PTPN6 (SHP1) | Tyrosine-Protein Phosphatase Non-Receptor Type 6 | AMPKA2/CAMKK2/CDK2/INSR | PPIN/GWAS/COEXPRESSION |

Collectively, IREP consists of more than 20% (15 validated DAGs from the T2D-protomodule and 101 candidates) of gene products confidently associated with the T2D risk.

### The insulin receptor-containing endosome network (IREN) displays a type-2 diabetes disease module architecture

A disease module can be defined as a connected subnetwork showing mechanistic evidence for a phenotype [16, 17]. To identify the molecular mechanisms associated with IREP, we constructed a PPIN of IR-containing endosomes. The cytoHubba algorithm was used to compute and to rank nodes in the network [37]. The resulting collated PPIN is formed by 313 nodes and 1147 edges (55% of IREP proteins; named IREN, Insulin Receptor Endosome Network). The general topology of IREN is based on few major hubs, with the kinase Cdk2 displaying the highest centrality. Relatively large nodes represented by the IR itself, proteins of the actin cytoskeleton (ACTB), and those involved in vesicular trafficking (CAV1) were observed. More peripheral nodes were also present as follows: coats (GOLGA2, CLTC), V-ATPase subunits (ATP6V1A), and cargos (APOA1) (Fig 3 and S5Table, sheet IREN).

**Figure 3.**
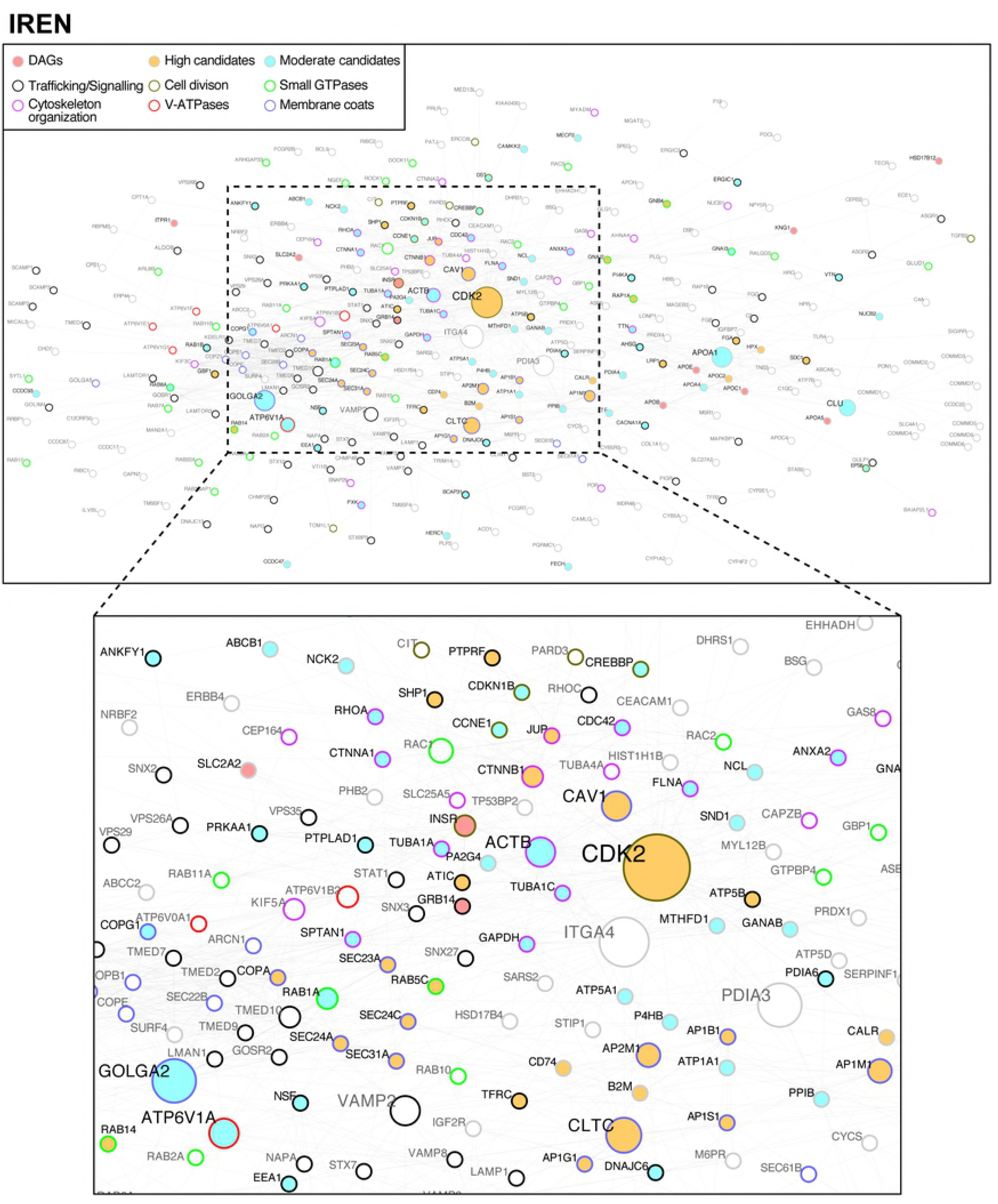
The physical protein interaction network of IR-containing endosomes (IREN) has a Cdk2 centrality and is highly associated with type 2 diabetes risk. The 557 IREP proteins (Table S4-sheet IREP-HUGO) were grouped and linked according to their physical association. The resulting network is formed by 313 nodes and 1147 edges (56% of IREP proteins). The general topology of IREN is based on few major hubs, with the kinase Cdk2 displaying the highest centrality (Table S5). Candidates (yellow and blue colors and black characters; Tables 1 and 2) and DAGs (pink color and black characters) form a single-connected disease module of 94 nodes (33% of IREN nodes) with 330 interactions (28,7% of IREN interactions). An expansion to the first level of adjacent nodes results in a connected subnetwork of 272 nodes (88% of nodes) covering 92% of interactions (1070 out of 1147 IREN interactions; Figures 3 and S3). The functional groups are represented according to the colors of the borders indicated in the legends.

From the 101 high-confidence candidates (Tables 1 and S9Table) and 15 of the 184 validated DAGs identified in IREP, 94 of the candidates and 10 of the DAGs are present in IREN. They form a single-connected subnetwork of 94 nodes with 330 interactions (Fig 3 and S5Table). To test whether this module could arise by chance in the context of IR endocytosis, we made random reiterations of any 94 nodes of IREN. The results showed that the subnetwork is robust (empirical p-value < 10^−4^; Fig S2). Its collective influence was analyzed by expanding it to the first adjacent nodes. This resulted in a connected network of 271 nodes (88% of nodes) and 1070 out of 1147 IREN interactions (Figs 3 and S3), coverage that is largely more than expected by chance (p-value < 10^−4^, Fig S2). GO analysis also revealed an enrichment for vesicle transport (p< 1.85 < 10-^51^) and response to oxygen species (p < 8.52 × 10^−20^), with the most enriched cell components being endosomes (p < 7.00 × 10^−25^), Golgi apparatus (p < 1.77 × 10-^27^) and endoplasmic reticulum (p < 2.13 × 10^−23^), showing that the T2D-protomodule expansion coincides with a functional expansion (Fig 3 and S5Table, sheet IREN, GO analysis). Of the 94 of the 101 high-confidence candidates in IREP, 94 have credible tyrosine phosphorylation motifs with the IREN kinases (Table1 and S9, S10 Tables). Of the 87 present in IREN, 71 have at least one of their kinase-substrate interactions confirmed in IREN (Fig 3 and S10Table), further emphasizing the mechanistic association. Taken together, these results indicate that IREN is a T2D-disease module.

### Candidate hub Cdk2 regulates the association of IR with microtubules

A prerequisite in the description of interaction maps is the validation of hubs in terms of perturbation responses. We then tested examples here of hub complexes in term of insulin response. We verified first whether Cdk2, which displays the highest centrality and is a high-confidence candidate (Table 1 and S1AFig), is indeed associated with key elements. Microtubules, for instance, rely on dimerization of tubulin subunits alpha and beta for their assembly and the association of internalized cargos to microtubules has been associated to late events of trafficking. We noticed that the tubulin alpha subunit (TUBA), a reported substrate for the IR [38, 39], is preponderant within the IREN (Fig 3). We show that TUBA indeed readily associates with the IR after insulin stimulation in HEK293 cells, while TUBB has a different profile (Fig 4A). In addition, the association was nearly abolished upon Cdk2 overexpression, confirming the presence of complexes and that Cdk2 presence has the capacity to regulate interactions within IREN (Fig 4A).

**Figure 4.**
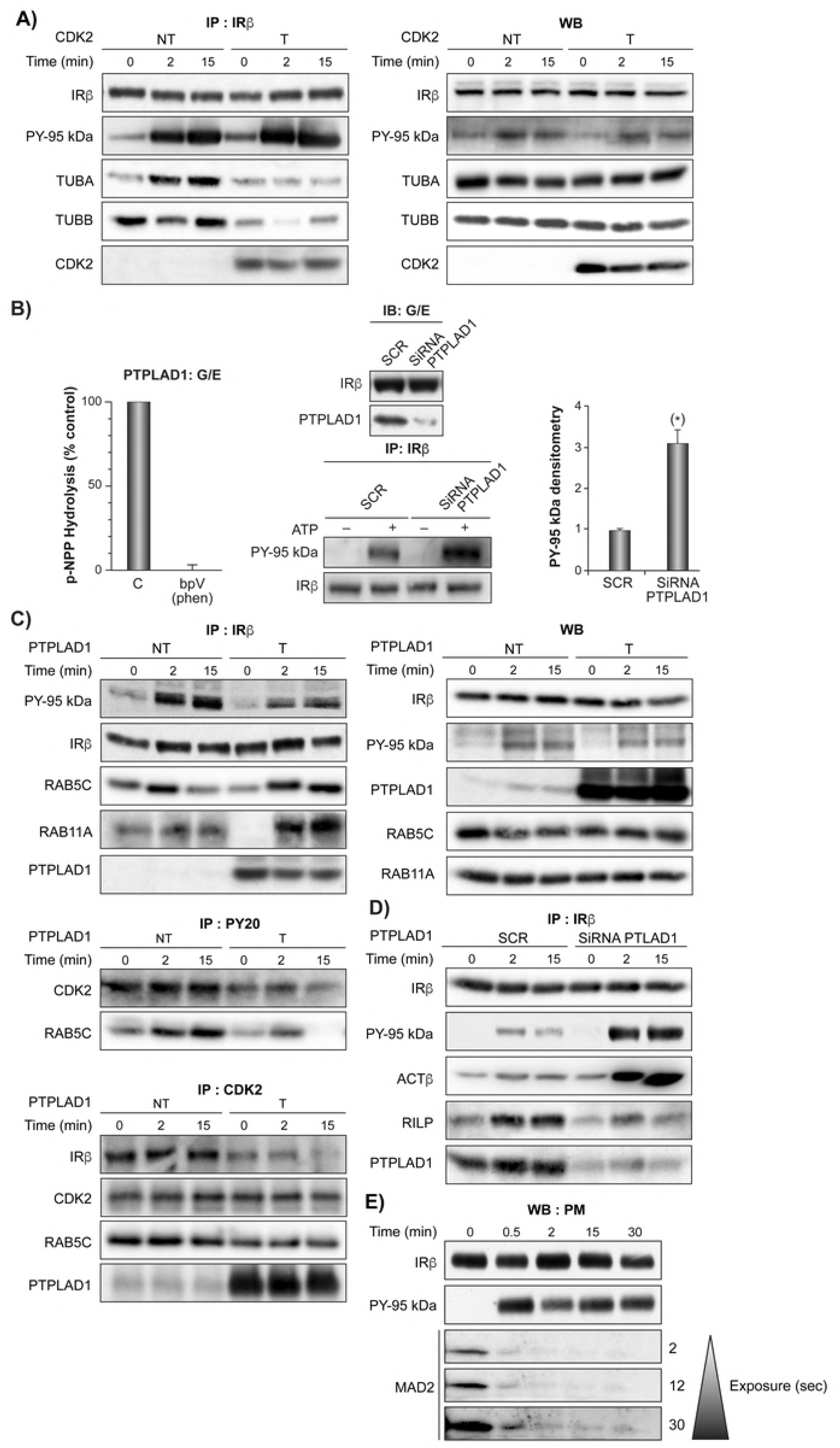
Cdk2 and PTPLAD1 interact with IR complex organization. (A) HEK293 cells were transfected with pcDNA3-CDK2 (T) or pcDNA3 (NT) for 48 hours. They were preincubated in serum-free medium for 5 hours and then stimulated for the indicated times with insulin (35 nM). Left panel: IR immunoprecipitation (IP: IRb), IR autophosphorylation (PY 95 kDa) and CDK2, TUBA and TUBB presence. Right panel: Immunoblots (WB) of CDK2, IR-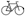-subunit, TUBA, B (pieces of the same membrane except PY-95 kDa (PY20 antibody); 3 independent experiments). (B) IR autophosphorylation increases in isolated endosomes depleted of PTPLAD1. Right panel: Rats were injected with a scrambled (SCR) or siRNA oligonucleotide targeting PTPLAD1 for 48 hours. The G/E fractions were then prepared from livers at their IR concentration time-peak (2 minutes after insulin injection; 1.5 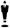 g/100 g, b.w.). The presence of IR and PTPLAD1 was verified by immunoblot (IB: G/E, input 50 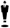 g of protein, pieces of the same membrane). IR immunoprecipitation (IP: IR 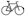) and IR autophosphorylation (PY 95 kDa) were measured after suspending endosomes in a cell-free system in the presence of ATP for 2 minutes at 37 ^o^C. After stopping the reaction, autophosphorylation was detected by immunoblotting using an anti-phosphotyrosine antibody (PY20). Normalized values shown in the right panel are means ± s.d. (* P<0.001 n=3). Left panel: PTPLAD1 was immunoprecipitated from the same fractions (input 30 mg protein of solubilized G/E) and incubated with p-NPP in the presence or absence of 50 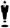 M bpV(phen). The measured activity was expressed as a percentage of 0.55 +/- 0.8 mmoles/min/mg of cell extract, n=4. (C) Cells were transfected with PTPLAD1-pcDNA3 (T) or pcDNA3 (NT) for 48 hours, incubated in serum-free medium for 5 hours and then stimulated for the indicated times with insulin (35 nM). The panel on the right shows immunoblots of PTPLAD1, IR-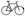 subunit, Rab5a and Rab11c from the total cell lysates. Left panel, IPs of IR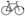, phosphotyrosine (PY20 antibody, middle left), and Cdk2 (bottom left) (3 independent experiments). (D) PTPLAD1 siRNA knockdown. IPs of the IR 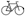 subunits were resolved by SDS-PAGE and were blotted for the indicated proteins (3 independent experiments). (E) The plasmamembrane (PM) fractions were prepared from rat liver at the indicated time following the injection of insulin (1.5 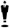 g/100 g b.w.). Fractions were monitored for the PM-associated MAD2 by immunoblotting (50 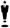 g of protein).

### PTPLAD1 expression regulates the IR autophosphorylation activity and association with Cdk2, Rab5c, Rab11a and actin

Compared with Cdk2, PTPLAD1 is an example of good centrality, but it is poorly studied and identified as a moderate candidate (S9Table and Fig 3). Because the fatty acid elongation enzymatic activity was not confirmed, it was recently hypothesized that PTPLAD1 (HACD3) is involved in the elongation of specialized forms of 3-OH acyl-CoAs, such as those containing a short or branched alkylic chain [40]. An interaction with Rac1 was also reported [41]. PTPLAD1 has a well-positioned conserved cysteine C(X)5K motif in the soluble cytosolic loop, residues 257-279, and its partial deletion in cultured HEK293 cells coincided with IR tyrosine hyper-phosphorylation [15]. We verified whether PTPLAD1 acts on IR tyrosine phosphorylation outside a whole-cell context. We used an in vitro assay, whereby IR-loaded endosomes were incubated in the presence of ATP. We observed that a prior siRNA-mediated depletion in rat PTPLAD1 nearly abolished the PTPLAD1 presence from isolated endosomes, which coincided with a marked increase in IR autophosphorylation, demonstrating that the IR tyrosine-phosphorylated state is modified by PTPLAD1 (Fig 4B right panels). Low, but consistent, enzymatic activity towards the artificial substrate pNPP was measured (Fig 4B left panel) resembling the loss of PTP activity towards the IR observed previously observed after membrane solubilization [42]. In accordance with these results, and with prior PTPLAD1 depletion experiments [15], we observed that overexpression of PTPLAD1 in HEK293 cells markedly decreases the IR tyrosine-phosphorylated state (Fig 4C). Of further interest, the candidate Rab5c (Table 1) also associates with the IR and this association increases in an insulin-regulated manner upon PTPLAD1 overexpression (Fig 4C). Rab5a and b are well documented as playing a role in the early events of EGFR endocytosis but the role of Rab5c remains unclear [43]. Rab5c may therefore be particularly important for IR action as we noted the presence of an IR phosphorylation motifs located in the GTP binding site (Y83) (S10Table) that resembles an inhibitory feedback loop described previously for Rab24 [44]. In support of this finding, we detected Rab5c and Cdk2, but not Rab11a, in anti-phosphotyrosine affinity complexes, and this association was abolished after PTPLAD1 overexpression (Fig 4C). In addition, both IR and Rab5c were present in Cdk2 affinity complexes, and this association decreased following PTPLAD1 overexpression (Fig 4C). To further test the importance of PTPLAD1 on IR routing, we verified and noticed an insulin-dependent association of IR with Rab11a, a known marker of endosomal recycling [22]. This supports a role for PTPLAD1 in cycling from endosomes to the plasmamembrane (PM). On another hand, we confirmed that under the same circumstances PTPLAD1 deletion, using siRNA, increases IR tyrosine phosphorylation and the presence of actin in IR immunoprecipitates (Fig 4D) [15].

We observed an insulin-dependent recruitment of the Rab- interacting lysosomal protein (RILP) in IR immunoprecipitates that was nearly abolished by PTPLAD1 deletion (Fig 4D). RILP was demonstrated to be required for EGFR confinement and degradation in late endosome compartments [45] and is an inhibitor of V-ATPase activity [46]. This supports the idea of a key role for PTPLAD1 in early events of IR internalisation and recycling via RAB5c, Rab11a and actin cytoskeleton elements. Collectively, the data support the presence of dynamic insulin-dependent interactions for Cdk2 between the IR, PTPLAD1, Rab5c, Rab11a, tubulin and actin cytoskeletons whereby PTPLAD1 controls IR tyrosine phosphorylation and sorting.

To verify the idea that cell cycle components have expanded their action on endocytic traffic, we verified whether the protein MAD2, which binds to the MAD2-interacting motif (MIM) located in the carboxyterminal domain of the IR β-subunit during clathrin-mediated endocytosis [47], is responsive to insulin at the cell surface. The results demonstrate that MAD2 readily disappears from the PM fractions following IR tyrosine kinase activation (Fig 4E), thus supporting the idea that cells use cell cycle regulators for both early [47] and later events of IR endocytosis.

### Inhibition of V-ATPase shifts the IR accumulation in endosomes in vivo

V-ATPase subunits are well represented in IREP (Fig 1 and S1Table), forming large, more peripheral nodes in IREN (Fig 3) with ATP6V1A being identified here as moderate candidate (S9Table) for type 2 diabetes risk. It was previously demonstrated that V-ATPase inhibition decreases IR recycling to the plasma membrane in cultured hepatocytes [48]. We thus verified whether the kinetics of IR endocytosis are affected in vivo after treatments with two different potent V-ATPases inhibitors. We observed that the peak of IR accumulation in endosomes is markedly shifted towards later stages of endocytosis following either concanamycin A or bafilomycin A1 pretreatments as demonstrated by immunoblotting and hexokinase activity measurements (Fig 5A). Concanamycin A does not affect IR tyrosine phosphorylation in vitro, suggesting that V-ATPase inhibitors do not inadvertently function through PTPs inhibition (Fig 5B left panel). We noted however a strong and consistent threonine phosphorylation signal that was readily abolished by concanamycin A (Fig 5B right panel), suggesting the presence of additional feedback loop layers, which have yet to be characterized, informing the cell that the lumenal acidification process is optimized. We verified whether V-ATPases elements contains IR phosphorylation motifs. The kinase network analysis indicated that ATP6V1A (S9Table) and ATP6V1E1 are indeed strong candidate substrates for the IR as well as for Cdk2 and AMPK (PRKAA1) (Fig 5C and S10Table).

**Figure 5.**
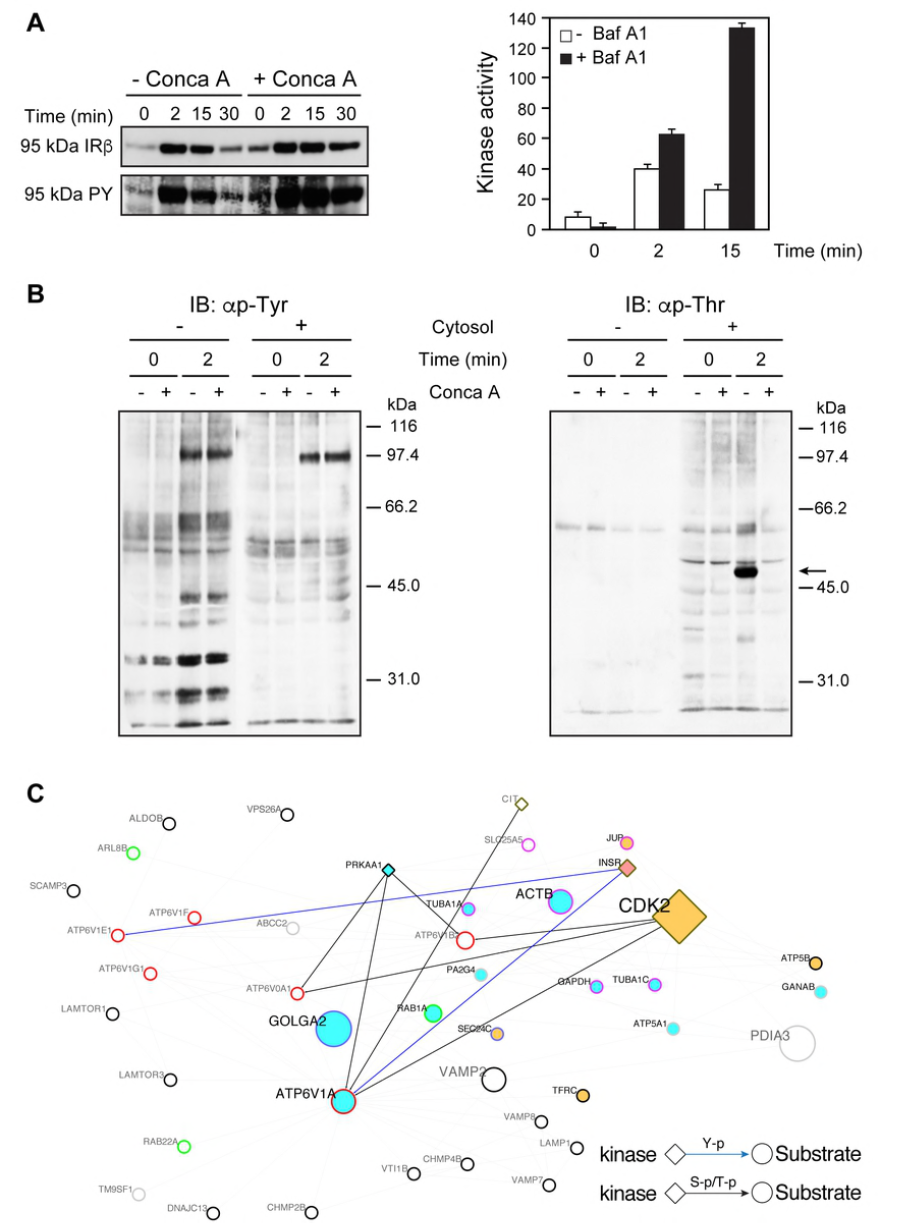
The pharmacological inhibition of the V-ATPase hub activity delays the time peak of IR accumulation in endosomes. Rats that were treated with concanamycin A (Conca A, 4.0 µg/100 g, b.w.) or were left untreated, were then stimulated with insulin (1.5 µg/100 g, b.w.) for the indicated time and the G/E fractions were isolated. (A) Left panel, immunoblot of IR using the anti-IRβ subunit or aPY20 (95 kDa PY) antibodies (50 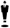 g of protein). Right panel, rats were left untreated or treated with bafilomycin A1 (Baf A1, 0.5 µg/100 g, b.w.). IRs from G/E fractions prepared at the noted time following insulin administration (1.5 µg/100 g, body weight) were partially purified by WGA-sepharose affinity chromatography and subjected to exogenous kinase assay. ^32^P incorporation into poly Glu-Tyr (4 :1) is expressed as pmol/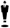 g protein. Values shown are means ± s.d. (P<0.0001, 2 minutes and 15 minutes, n=3). (B) G/E liver fractions were prepared at their IR concentration time peak (2 minutes after insulin injection; 1.5 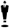 g/100 g b.w.) and immediately suspended in the cell-free system for 0 and 2 minutes at 37 ^o^C and in the presence of ATP and the absence or presence of fresh cytosol (diluted 1/10) and Conca A. After stopping the reaction (0 and 2 minutes), the fractions were immunoblotted (input 50 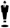 g of protein; 12 % resolving gels) with the anti-phosphotyrosine (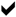p-Tyr, left panel) or anti-phosphothreonine (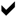p-Thr, right panel) antibodies. (C) Subnetwork extracted from IREN (Figure 3) depicting the connectivity of V-ATPase subunits. The V-ATPase subunits ATP6V1A, ATP6V1E1, ATP6VDA1 and ATP6V1B2 containing high confidence IR-tyrosine kinase phosphorylation and Ser/Thr kinases Cdk2, PRKAA1 (AMPK) and Citron phosphorylation motifs (Table S10) are marked according to the legend.

## Discussion

Using a combination of cell fractionation and computational approaches, we found a T2D disease module in IR-containing endosomes. The starting point of our analysis was a list of seed genes with established genetic T2D association and high GWAS p-values (1 × 10^−8^) against the background of random variation. They carry enough information to build a robust T2D-protomodule (Fig 2). The functional specialization of the T2D-protomodule also found in IREN (Fig 3) is in accord with the connection of these processes (protein transport, transcriptional factors and response to oxygen) in insulin action [1, 2]. The topological features of a scale-free network, with the view that hubs with the highest influence represent important points in biological networks [16, 37], coupled with the large enrichment in T2D genetic risk is particularly well represented by Cdk2 (Fig 3, Table 1). Cdk2 regulators were independently and repeatedly reported by GWAS and their role, with many other common variants, was interpreted more in terms of insulin production and secretion indicating that the beta-cell is a more appropriate place to find a T2D-disease module [11, 13, 49, 50]. Indeed, mice lacking Cdk2 are viable [51, 52] and targeted Cdk2 deletion in the pancreas induces glucose intolerance primarily by affecting glucose-stimulated insulin secretion [53]. Similar to endosomes, the secretory pathway consists of multiple dynamic compartments linked via anterograde and retrograde transport [54, 55]. The T2D-disease module thus can be co-functional in endosomes and insulin-secreting cells. In this regard, in the liver the presence of insulin-regulated Cdk2/cyclinE/p27^kip1^complexes having a capacity to inhibit hybrid endosome formation in vitro has been previously reported [56].

In contrast with Cdk2, PTPLAD1 has less topological influence in IREN and is a less-studied protein. PTPLAD1 is, however, functionally well connected, as the control of IR activity may be achieved at several endosomal targets by PTPLAD1 that, together with Cdk2, seems to have a considerable local influence on actin and microtubule networks, Rabs and V-ATPase. The finding that IR complexes are under the control of PTPLAD1 would also be particularly important, because PTPLAD1 mobilization in response to insulin inputs has also interesting consequences by favoring tyrosine phosphorylated-IR quanta formation, which is considered as an emergent property of endosomes as signaling devices [5]. This PTP activity is yet to be fully characterized and can be supported elsewhere in the cell by the small fraction of the endoplasmic reticulum-associated PTP-1B with high specific activity that can reach the plasmamembrane at specific points of cell-cell contact [57], by cytosolic PTPs (SHP1/2) (Fig 2, Table 1) that couple to RTK phosphorylation in a negative feedback manner at the PM with longer delays [58], and by PTPRs that are thought to display low specific activity towards basal RTK autophosphorylation activities occurring at the cell surface [59]. Through a concerted action on microtubules and actin elements IREN supports a model in which Cdk2 controls the microtubules-based traffic, and PTPLAD1 is an insulin-dependent switch deciding the choice of IR interaction with microtubules versus actin routing events (Fig 4). Interesting times are ahead for investigating insulin responses in the context of IREN. The question arises as to the extent of crosstalk between the IR-Tyr kinase and the predominantly Ser**/**Thr kinases (Cdk2, AMPK) that drive IR trafficking and signaling, and when and where this crosstalk occurs. Apart from the presence of multiple high-confidence substrates for Cdk2 and AMPK in IREN, the current results strongly point to the internalized IR as a relatively pleiotropic *writer* in the disease module (Tables 1 and S9, S10 Tables; For example, ATP6V1E1: Y-464; AMPK: Y-247; ATIC: Y-151; Rab5c: Y-83) and PTPLAD1 as the insulin-dependent *eraser* with short delay. A related challenge will be systematically matching these phosphor-sites to their cognate physiological *readers* [60].

Another connected example of the IR regulatory mechanism associated with the T2D genetic risk concerns the marked effect of the proton pumping activity on IR trafficking in vivo (Fig 5). A concrete problem for the cell concerns the energy sources, and it seems that an efficient solution was found to connect IR activity with intermediary metabolism and trafficking by linking V-ATPase subunits (continuous energy demand) with AMPK (energy sensor and action on IR trafficking) and the metabolic enzyme ATIC (ATP production) (Fig 5) further supporting the idea of the presence of an IR/ATIC/AMPK/PTPLAD1 circuit [15, 61]. The decreased presence of ATIC homodimers, using a small interface interactor, indeed activates AMPK and improve glucose intolerance in a mouse model [62]. We also noted the presence of related candidate enzyme, MTHFD1 (Fig 3 and S9Table: PPIN, GWAS, co-expression). The fact that V-ATPase controls the activity of AMPK [63] emphasizes the idea that all the conditions are present in IREN to auto-regulate this node and thus IR routing, signaling and hepatic clearance in relation to global cell energy status. The presence of the V-ATPase inhibitor RILP [46] in IR immunoprecipitates, which was nearly abolished by PTPLAD1 deletion (Fig 4D), further supports the idea that PTPLAD1 has a large capacity for action to decide IR routing towards early versus late compartments [45].

A facet of IR trafficking in endosomes that can affect indirectly insulin production and secretion is the insulin dissociation/degradation sequence in endosomes, which supports efficient hepatic insulin clearance [3, 32]. A reduction in hepatic insulin clearance is viewed as an adaptive mechanism that relieves the burden on pancreatic beta-cells [8, 64]. On the other hand, as shown by a mouse model, moderate chronic hyperinsulinemia can be the primary mechanism resulting in insulin resistance [65]. The idea that the complex genetic heterogeneity converges towards a single module co-functional in insulin-producing and target cells, implies a mechanistic promiscuity between insulin signaling, transport and production, that can explain the prevalence of insulin clearance in insulin sensitivity found in some animal models [66].

We acknowledge that some endosomal structures might not be accessible to the IR β-subunit antibody and the limitations inherent to the fractionation approaches such as true tubular connection between different organelles versus contaminants [19, 67]. Nonetheless, the present IREN helps us narrow the search space of the full organism interactome and focus a search in a well-localized network neighborhood. Quantitative proteomic approaches are needed to establish how changes in endosomes occur in space and time according to low (around 10 % saturating) versus large saturating insulin doses (50 % saturating here). This will provide a more complete picture of IREN dynamics that takes the in vivo polarized situation into account [7].

In conclusion, our results establish that the endosomal apparatus contains a T2D disease module located in close proximity to the IR. It senses the state of IR activation and seems co-functional with insulin secretion and islets biology. It helps to explain disease heterogeneity and represents a valuable new resource to understand insulin action and to classify related metabolic traits. Rewiring a network, distorted under the combined genetic and environmental pressures, with designed surface interactors [33], provides a mechanistic rationale for the exploration of personalized medicine and elaborate new necessary drugs [1, 2, 68].

## Methods

### Cell Fractions

Harlan Sprague-Dawley rats (female 120-140 g, b.w.) were purchased from Charles River Ltd. (St. Constant, Québec, Canada) and were maintained under standard laboratory conditions with food and water available *ad libitum*, except that the food was removed 18 hours before the experiments. All animal procedures were approved by the CPA-CRCHUQ (certificate 055-3). The G/E and the PM fractions were prepared and characterized in terms of enzyme markers, electron microscopy (EM) and ligand-mediated endocytosis, as originally described [3]. The G/E fraction was also characterized in terms of proteomic survey and construction of the protein interaction network (GEN) [15]. IR-immuno-enriched endosomes were prepared as originally depicted [19], starting from the mixed hepatic Golgi/endosomal (G/E) fraction with only minor modifications [20]. Dynabeads (Dynal-A, Invitrogen, San Francisco, CA, USA) that were pretreated with 0.1% BSA and coated with the anti-IR β-subunit antibody were incubated with freshly prepared G/E fractions (10 mg of protein) for 1 hour at 4°C under gentle agitation. Beads were then rapidly rinsed before being subjected to EM, immunoblotting and mass spectrometry (MS) analysis. There was no major differences in the size and morphology of the vesicles immuno-isolated after 2 minutes or after 15 minutes of stimulation. They were relatively homogeneous with a diameter of 70-200 nm and some tubular elements.

### Protein in-gel digestion

Beads were washed 3 times with 50 mM ammonium bicarbonate buffer. They were suspended in 25 µl of 50 mM ammonium bicarbonate, following which trypsin (1 µg) was added. Proteolysis was done at 37°C and stopped by acidification with 3% acetonitrile-1% TFA-0.5% acetic acid. Beads were removed by centrifugation, and peptides were purified from the supernatant by stage tip (C18) and vacuum dried before MS injection. Samples were solubilized into 10 µl of 0.1% formic acid and 5 µl was analyzed by mass spectrometry [69].

### Mass spectrometry

Peptide samples were separated by online reverse-phase (RP) nanoscale capillary liquid chromatography (nanoLC) and analyzed by electrospray mass spectrometry (ES MS/MS). The experiments were performed with an Agilent 1200 nano pump connected to a 5600 mass spectrometer (AB Sciex, Framingham, MA, USA) equipped with a Nanoelectrospray ion source. Peptide separation occurred on a self-packed PicoFrit column (New Objective, Woburn, MA) packed with Jupiter (Phenomenex, Torrance, CA) 5 ul, 300A C18, 15 cm x 0.075 mm internal diameter. Peptides were eluted with a linear gradient from 2-30% solvent B (acetonitrile, 0.1% formic acid) in 30 minutes at 300 nl/min. Mass spectra were acquired using a data-dependent acquisition mode using Analyst software version 1.6. Each full scan mass spectrum (400 m/z to 1250 m/z) was followed by collision-induced dissociation of the twenty most intense ions. Dynamic exclusion was set for a period of 3 sec and a tolerance of 100 ppm. All MS/MS peak lists (MGF files) were generated using Protein Pilot (AB Sciex, Framingham, MA, USA, Version 4.5) with the paragon algorithm. MGF sample files were then analyzed using Mascot (Matrix Science, London, UK; version 2.4.0). MGF peak list files were created using Protein Pilot version 4.5 software (ABSciex) utilizing the Paragon and Progroup algorithms. (Shilov). MGF sample files were then analyzed using Mascot (Matrix Science, version 2.4.0) [70], and rodent databases (S1Table). The number of newly identified proteins plateaued at approximately 10-20% of total for the second and third experiments, indicating that we were close to the completion point with this method [71] (S3Fig).

### Databases and network analyses

Conversion to human orthologs was performed using the InParanoid8 database. The PPIN was generated from a listing of protein-coding genes generated and named according to HUGO database nomenclature. Proteins found to be associated with IR in hepatic endosomes were included in the analysis: ATIC, PTPLAD1, SHP1, CDK2, PLVAP1, CDKN1B and CCNE1. The interactions were curated using Y2H binary interactions of the CCSB human interactome, physical complexes and direct interactions from Intact, Database of Interacting Protein (DIP, UCLA), REACTOME, HITPREDICT and HINT databases, affinity complexes from BIOGRID and HPRD databases. Proteins having nonspecific interactions such as chaperones, ribosomal (RPL family) proteins, ubiquitylation and sumoylation processes (UBC, CUL), elongation factors were removed [72, 73]. The Cytoscape platform (Version 3.2.0) was used for network visualization [34]. Self-loops and duplicated edges were removed prior the analyses. The cytoHubba algorithm was used to compute and rank nodes according to their centrality « Betweenness and Connectivity » scores in the network [37, 72, 74] (S5Table). Cellular component grouping and functional analysis were performed after a gene ontology analysis with the Biological Networks Gene Ontology tool *(BINGO version 2.44).* Kinase predictions were performed with GPS 3.0 [75], phosphosites [76] and NetworKIN [77] version 3.0 (KinomeXplorer) using the high-throughput workflow option. Data from GPS 3.0 were additionally filtered by a differential score (difference between Score and Cut of) higher or equal to 1.0. Networking associations were considered if the Networkin score was observed to be higher than 2.0. Analyses were performed on November 10 2017.

### Candidate gene analysis and identification

#### GO analysis

We verified the probability of intracellular colocalization for candidates and seeds using the plugin BINGO adapted for the Cytoscape platform. We clustered the hybrid network (S4Figure) based on enrichment in the same cellular compartment by GO. In IREP proteins coming from Golgi-endosomal fractions, 21 seeds were found to be enriched in the Golgi apparatus (p < 5.6822 × 10^−14^, after correction) and endosomes (p < 1.1315 × 10^−16^, after correction). Of the 126 IREP candidates identified by PPIN, 32 have at least three interactors among the 21 Golgi-endosomal seeds. The analysis was expanded to other compartments with 7 candidates interacting each with three seeds in the cytosol cluster (p < 6.8728 × 10^−17^, after correction), 10 candidates in the endoplasmic reticulum (p < 1.6173 × 10^−12^, after correction), 25 in the plasmamembrane (p < 5.6570 × 10^−8^, after correction), and 13 in the extracellular region (p < 4.6024 10 × 10^−5^, after correction). Taken together, 54 nonredundant IREP coding genes among the 126 identified by PPIN were found to be colocalized with validated seeds based on GO analysis (S5Fig and S6Table).

#### Fine-mapping approach

We performed a linkage disequilibrium (LD) analysis and identified proximal SNPs correlated to diabetes GWAS signals (p ≤ 10^−3^) using replicated data as displayed in tables from the Wellcome Trust Case Control Consortium (WTCCC), GWAS Central portal, GWAS catalog or DIAGRAM GWAS-Metabochip or trans-ethnic data. This analysis provided a list of 130 IREP coding genes falling in genomic loci reliably associated with diabetes (S7Table).

#### Genes expression analysis

Most of the SNPs identified by GWAS are intergenic or fall in intronic regions of genes suggesting a regulatory role [9, 11]. Among the 130 candidates identified by fine-mapping, we verified which ones had SNPs experimentally shown to affect gene expression and to likely regulate some transcription factor binding as described in category-1 of high-confidence associations in the RegulomeDB database [78]. We identified 15 IREP coding genes fulfilling these criteria, consequently forming a first pool of IREP candidates based on gene expression regulation (Table S8). A second pool was made-up of IREP genes showing or predicted to have similar patterns of expression with at least three of the 184 seeds by RNA-Seq analysis and simultaneously sharing regulatory binding motifs either for transcription factors or for miRNA. The candidates and seeds pairs were considered coexpressed if they were mutually ranked among the top 1% of coexpressed genes pairs by the Genefriends database [79]. The transcription factor targets (TFTs) or microRNA targets were analyzed using the top 10 grouping of the Gene Set Enrichment Analysis [80, 81] with (p < 4.35 × 10-16 after correction for TFTs and p < 2.88 × 10-6 for miRNA targets). In all, 296 IREP coding genes were found to share TFTs with at least three diabetes genes compared with 112 for miRNA targets and 109 for RNA-Seq. Only 80 genes from the RNA-Seq analysis were considered for the second pool of candidates because they simultaneously showed some shared binding targets with at least three DAGs for TFs (72 genes) and/or for miRNA (28 genes). Taken together, 94 nonredundant IREP coding genes from the first and second pools are considered candidates based on shared regulatory elements with validated DAGs (S8Table).

#### IR endosomal autophosphorylation

IR endosomal autophosphorylation was measured as previously reported [42] with minor modifications [15]. *SiRNA* in vivo: Rats were injected via the jugular vein with a scrambled or predesigned stabilized rat PTPLAD1 sequence (100 mg/100 g bw; IVORY in vivo siRNA GGGGCAGUCUAAUUCGGUGUGCU, D-00203-0200-V; purified/desalted by RP/IEX-HPLC; Riboxx Life Sciences, Germany; Liver *In vivo* transfection reagent 5061, Altogen Biosystems, Las Vegas, CA) 48 and 24 hours before isolating the G/E fraction. The PTPLAD1 mRNA expression level was measured against GAPDH in liver sections using quantitative polymerase chain reaction (qPCR) and was decreased by 52 +/- 6.2%, n=3.

#### Cell culture and analysis

HEK293 cells were maintained in DMEM high-glucose medium with 10% foetal bovine serum. PTPLAD1 siRNA knockdown was performed as previously described [15] using the predesigned human sequence as follows: GACCCAGAGGCAGGUAAACAUUACA NM_016395_STEALTH_367. Cells were transfected using Lipofectamine 2000™ (Life Technologies) for 48 hours and subjected to the described experiments. For overexpression experiments PTPLAD1 WT and Cdk2 WT were cloned into the pcDNA3 expression vector. Transfection was performed with Lipofectamine 2000™ and plasmid DNA (300 ng/ml). Cells were preincubated at 37 °C without serum for 5 hours before insulin (35 nM) stimulation for the indicated times. Immunoprecipitation (IP) were done under solubilization conditions that preserve the integrity of insulin-dependent complexes (Empigen BB 0.3%, 2 hours, 4°C) [20].

#### Reagents and antibodies

Porcine insulin (I5523) was obtained from Sigma-Aldrich (St. Louis, MO, USA). The following antibodies were used: anti-phosphotyrosine (PY20, Sigma-Aldrich, St. Louis, MO, USA). The IR β-subunit (Sc-711), Rab5c (sc-365667) and Cdk2 (sc-163, sc-163AC) antibodies were obtained from Santa Cruz Biotechnology (Santa Cruz, CA, USA). The anti-PTPLAD1 was from Abcam (ab57143, Cambridge MA, USA). The anti-tubulin antibodies were obtained from Sigma-Aldrich (T5168, TUB 2.1, St. Louis, MO, USA). The anti-MAD2 was from Bethyl Laboratories (Montgomery, TX, USA). The RILP antibody was from Invitrogen (PA5-34357, Waltham, MA, USA). The generic anti-phosphothreonine was from Zymed (San Francisco, CA, USA). The antibody against Rab11a was from ThermoFisher Scientific (Rockford, IL, USA). Peroxidase-conjugated secondary antibodies were used (1:10,000, Jackson Immuno Research Laboratories, West Grove, PA, USA). Membranes (PVDF) were analyzed using a chemiluminescence kit (ECL, Perkin Elmer Life science, Boston, MA) or using an ImageQuant LAS 40 000 imager (GE Healthcare Biosciences, Baie d’Urfé, QC, CA). [γ-^32^P]-ATP (1000-3000 Ci/mmol) was from New England Nuclear Radiochemicals (Lachine, Québec). Other chemicals and reagents were of analytical grade and were purchased from Fisher Scientific (Sainte-Foy, Québec, CAN) or from Roche Laboratories (Laval, Québec, CAN).

## Acknowledgements

We thank Dr Christian R. Landry (Université Laval) for critical comments.

## Author contributions

Conceptualization, M. B-D, R. F.; Methodology, M. B-D, P. B., S. B., A. D., S. F., R. F.; Investigation, M.B -D, N. B., S. B., S. F., S. E.; Writing- Original draft, M. B -D; Writing-review and editing, R.F., S. E.; Funding acquisition, R. F.

## Supplemental Informations

Figure S1 LD display (Haploview) of the *CDK2* and *HACD3* genes.

Figure S2 Random reiterations (10 000) simulations of 94 nodes subnetworks.

Figure S3 IREP: Number of newly identified proteins from one experiment to another (tryptic peptides).

Figure S4 Hybrid module (T2D-protomodule/IREP).

Figure S5 Extracted T2D subnetwork

Table S1. Proteome: proteins and spectra reports.

Table S2. Source list of T2D and associated traits (glucose intolerance, obesity) genes.

Table S3. Selected DAGs and validated seeds.

Table S4. Listing of IREP proteins orthology and networks.

Table S5. IREN and T2D-protomodule construction with Hubaa; GO analysis.

Table S6. Gene Ontology (GO) subcellular analysis.

Table S7. Fine mapping analysis: LD analysis of IREP coding genes and DAGs variants.

Table S8. TF motifs and coexpression analysis.

Table S9. Tables of candidates: 63 IREP coding genes are validated for association to diabetes traits by at least two out of four distinct approaches. (PPIN) protein-protein interactions network. (GO) Gene Ontology, Sub-cellular colocalization. (GWAS) fine-mapping. (COEXPRESSION) same expression pattern.

Table S10. Kinase-substrate analysis based on Phosphositeplus, Networkin and GPS 3.0 databases.

## Acknowledgements

The research program in the R. Faure laboratory was funded by the National Sciences and Engineering Research Council of Canada (NSERC: 155751) and the Fondation du CHU de Québec. M. B-D acknowledges funding from the Fondation du CHU de Québec and the CRCHUQ. S.E. holds a FRQS junior investigator salary award.

## References

1. Boucher J, Kleinridders A, Kahn CR. Insulin receptor signaling in normal and insulin-resistant states. Cold Spring Harb Perspect Biol. 2014;6(1). Epub 2014/01/05. doi: 6/1/a009191 [pii] 10.1101/cshperspect.a009191. PubMed PMID: 24384568; PubMed Central PMCID: PMC3941218.

2. Haeusler RA, McGraw TE, Accili D. Biochemical and cellular properties of insulin receptor signalling. Nat Rev Mol Cell Biol. 2017. doi: 10.1038/nrm.2017.89. PubMed PMID: 28974775.

3. Bergeron JJ, Di Guglielmo GM, Dahan S, Dominguez M, Posner BI. Spatial and Temporal Regulation of Receptor Tyrosine Kinase Activation and Intracellular Signal Transduction. Annu Rev Biochem. 2016. Epub 2016/03/30. doi: 10.1146/annurev-biochem-060815-014659. PubMed PMID: 27023845.

4. Goh LK, Sorkin A. Endocytosis of receptor tyrosine kinases. Cold Spring Harb Perspect Biol. 2013;5(5):a017459. Epub 2013/05/03. doi: 5/5/a017459 [pii] 10.1101/cshperspect.a017459. PubMed PMID: 23637288.

5. Villasenor R, Kalaidzidis Y, Zerial M. Signal processing by the endosomal system. Curr Opin Cell Biol. 2016;39:53–60. doi: 10.1016/j.ceb.2016.02.002. PubMed PMID: 26921695.

6. Bergeron JJ, Di Guglielmo GM, Dahan S, Dominguez M, Posner BI. Spatial and Temporal Regulation of Receptor Tyrosine Kinase Activation and Intracellular Signal Transduction. Annu Rev Biochem. 2016;85:573–97. Epub 2016/03/30. doi: 10.1146/annurev-biochem-060815-014659. PubMed PMID: 27023845.

7. Zeigerer A, Wuttke A, Marsico G, Seifert S, Kalaidzidis Y, Zerial M. Functional properties of hepatocytes in vitro are correlated with cell polarity maintenance. Exp Cell Res. 2016. doi: 10.1016/j.yexcr.2016.11.027. PubMed PMID: 27916608.

8. Samuel VT, Shulman GI. Mechanisms for insulin resistance: common threads and missing links. Cell. 2012;148(5):852–71. Epub 2012/03/06. doi: S0092-8674(12)00217-6 [pii] 10.1016/j.cell.2012.02.017. PubMed PMID: 22385956; PubMed Central PMCID: PMC3294420.

9. Prasad RB, Groop L. Genetics of type 2 diabetes-pitfalls and possibilities. Genes (Basel). 2015;6(1):87–123. doi: 10.3390/genes6010087. PubMed PMID: 25774817; PubMed Central PMCID: PMCPMC4377835.

10. Fuchsberger C, Flannick J, Teslovich TM, Mahajan A, Agarwala V, Gaulton KJ, et al. The genetic architecture of type 2 diabetes. Nature. 2016. Epub 2016/07/12. doi: nature18642 [pii] 10.1038/nature18642. PubMed PMID: 27398621.

11. Visscher PM, Wray NR, Zhang Q, Sklar P, McCarthy MI, Brown MA, et al. 10 Years of GWAS Discovery: Biology, Function, and Translation. Am J Hum Genet. 2017;101(1):5–22. doi: 10.1016/j.ajhg.2017.06.005. PubMed PMID: 28686856; PubMed Central PMCID: PMCPMC5501872.

12. Dimas AS, Lagou V, Barker A, Knowles JW, Magi R, Hivert MF, et al. Impact of type 2 diabetes susceptibility variants on quantitative glycemic traits reveals mechanistic heterogeneity. Diabetes. 2014;63(6):2158–71. doi: 10.2337/db13-0949. PubMed PMID: 24296717; PubMed Central PMCID: PMCPMC4030103.

13. Hannou SA, Wouters K, Paumelle R, Staels B. Functional genomics of the CDKN2A/B locus in cardiovascular and metabolic disease: what have we learned from GWASs? Trends Endocrinol Metab. 2015;26(4):176–84. Epub 2015/03/07. doi: S1043-2760(15)00023-5 [pii] 10.1016/j.tem.2015.01.008. PubMed PMID: 25744911.

14. Wood AR, Jonsson A, Jackson AU, Wang N, van Leewen N, Palmer ND, et al. A Genome-Wide Association Study of IVGTT-Based Measures of First-Phase Insulin Secretion Refines the Underlying Physiology of Type 2 Diabetes Variants. Diabetes. 2017;66(8):2296–309. doi: 10.2337/db16-1452. PubMed PMID: 28490609; PubMed Central PMCID: PMCPMC5521867.

15. Boutchueng-Djidjou M, Collard-Simard G, Fortier S, Hebert SS, Kelly I, Landry CR, et al. The Last Enzyme of the De Novo Purine Synthesis Pathway 5-aminoimidazole-4-carboxamide Ribonucleotide Formyltransferase/IMP Cyclohydrolase (ATIC) Plays a Central Role in Insulin Signaling and the Golgi/Endosomes Protein Network. Mol Cell Proteomics. 2015;14(4):1079–92. Epub 2015/02/18. doi: M114.047159 [pii] 10.1074/mcp.M114.047159. PubMed PMID: 25687571.

16. Barabasi AL, Gulbahce N, Loscalzo J. Network medicine: a network-based approach to human disease. Nat Rev Genet. 2011;12(1):56–68. Epub 2010/12/18. doi: nrg2918 [pii] 10.1038/nrg2918. PubMed PMID: 21164525; PubMed Central PMCID: PMC3140052.

17. Sharma A, Menche J, Huang CC, Ort T, Zhou X, Kitsak M, et al. A disease module in the interactome explains disease heterogeneity, drug response and captures novel pathways and genes in asthma. Hum Mol Genet. 2015;24(11):3005–20. doi: 10.1093/hmg/ddv001. PubMed PMID: 25586491; PubMed Central PMCID: PMCPMC4447811.

18. Posner BI, Bergeron JJ. Assessment of internalization and endosomal signaling: studies with insulin and EGF. Methods Enzymol. 2014;535:293–307. Epub 2014/01/01. doi: B978-0-12-397925-4.00017-1 [pii] 10.1016/B978-0-12-397925-4.00017-1. PubMed PMID: 24377930.

19. Jin M, Saucan L, Farquhar MG, Palade GE. Rab1a and multiple other Rab proteins are associated with the transcytotic pathway in rat liver. J Biol Chem. 1996;271(47):30105–13. Epub 1996/11/22. PubMed PMID: 8939959.

20. Fiset A, Xu E, Bergeron S, Marette A, Pelletier G, Siminovitch KA, et al. Compartmentalized CDK2 is connected with SHP-1 and beta-catenin and regulates insulin internalization. Cell Signal. 2011;23(5):911–9. Epub 2011/01/26. doi: S0898- 6568(11)00020-9 [pii] 10.1016/j.cellsig.2011.01.019. PubMed PMID: 21262353.

21. Rink J, Ghigo E, Kalaidzidis Y, Zerial M. Rab conversion as a mechanism of progression from early to late endosomes. Cell. 2005;122(5):735–49. Epub 2005/09/07. doi: S0092-8674(05)00697-5 [pii] 10.1016/j.cell.2005.06.043. PubMed PMID: 16143105.

22. Welz T, Wellbourne-Wood J, Kerkhoff E. Orchestration of cell surface proteins by Rab11. Trends Cell Biol. 2014;24(7):407–15. Epub 2014/03/29. doi: S0962- 8924(14)00033-6 [pii] 10.1016/j.tcb.2014.02.004. PubMed PMID: 24675420.

23. Andersen JN, Del Vecchio RL, Kannan N, Gergel J, Neuwald AF, Tonks NK. Computational analysis of protein tyrosine phosphatases: practical guide to bioinformatics and data resources. Methods. 2005;35(1):90–114. Epub 2004/12/14. doi: S1046- 2023(04)00175-6 [pii] 10.1016/j.ymeth.2004.07.012. PubMed PMID: 15588990.

24. Ramachandran C, Aebersold R, Tonks NK, Pot DA. Sequential dephosphorylation of a multiply phosphorylated insulin receptor peptide by protein tyrosine phosphatases. Biochemistry. 1992;31(17):4232–8. Epub 1992/05/05. PubMed PMID: 1373652.

25. Hashimoto N, Feener EP, Zhang WR, Goldstein BJ. Insulin receptor protein-tyrosine phosphatases. Leukocyte common antigen-related phosphatase rapidly deactivates the insulin receptor kinase by preferential dephosphorylation of the receptor regulatory domain. J Biol Chem. 1992;267(20):13811–4. Epub 1992/07/15. PubMed PMID: 1321126.

26. Moller NP, Moller KB, Lammers R, Kharitonenkov A, Hoppe E, Wiberg FC, et al. Selective down-regulation of the insulin receptor signal by protein-tyrosine phosphatases alpha and epsilon. J Biol Chem. 1995;270(39):23126–31. Epub 1995/09/29. PubMed PMID: 7559456.

27. Shintani T, Higashi S, Takeuchi Y, Gaudio E, Trapasso F, Fusco A, et al. The R3 receptor-like protein tyrosine phosphatase subfamily inhibits insulin signalling by dephosphorylating the insulin receptor at specific sites. J Biochem. 2015;158(3):235–43. Epub 2015/06/13. doi: mvv045 [pii] 10.1093/jb/mvv045. PubMed PMID: 26063811.

28. Gall WE, Higginbotham MA, Chen C, Ingram MF, Cyr DM, Graham TR. The auxilin-like phosphoprotein Swa2p is required for clathrin function in yeast. Curr Biol. 2000;10(21):1349–58. Epub 2000/11/21. doi: S0960-9822(00)00771-5 [pii]. PubMed PMID: 11084334.

29. Sousa R, Lafer EM. The role of molecular chaperones in clathrin mediated vesicular trafficking. Front Mol Biosci. 2015;2:26. Epub 2015/06/05. doi: 10.3389/fmolb.2015.00026. PubMed PMID: 26042225; PubMed Central PMCID: PMC4436892.

30. Dubois MJ, Bergeron S, Kim HJ, Dombrowski L, Perreault M, Fournes B, et al. The SHP-1 protein tyrosine phosphatase negatively modulates glucose homeostasis. Nat Med. 2006;12(5):549–56. PubMed PMID: 16617349.

31. Tuomikoski T, Felix MA, Doree M, Gruenberg J. Inhibition of endocytic vesicle fusion in vitro by the cell-cycle control protein kinase cdc2. Nature. 1989;342(6252):942–5.

32. Duckworth WC, Bennett RG, Hamel FG. Insulin degradation: progress and potential. Endocr Rev. 1998;19(5):608–24. Epub 1998/10/30. PubMed PMID: 9793760.

33. Sahni N, Yi S, Taipale M, Fuxman Bass JI, Coulombe-Huntington J, Yang F, et al. Widespread macromolecular interaction perturbations in human genetic disorders. Cell. 2015;161(3):647–60. Epub 2015/04/25. doi: S0092-8674(15)00430-4 [pii] 10.1016/j.cell.2015.04.013. PubMed PMID: 25910212; PubMed Central PMCID: PMC4441215.

34. Doncheva NT, Assenov Y, Domingues FS, Albrecht M. Topological analysis and interactive visualization of biological networks and protein structures. Nat Protoc. 2012;7(4):670–85. Epub 2012/03/17. doi: nprot.2012.004 [pii] 10.1038/nprot.2012.004. PubMed PMID: 22422314.

35. Oti M, Snel B, Huynen MA, Brunner HG. Predicting disease genes using protein-protein interactions. J Med Genet. 2006;43(8):691–8. Epub 2006/04/14. doi: jmg.2006.041376 [pii] 10.1136/jmg.2006.041376. PubMed PMID: 16611749; PubMed Central PMCID: PMC2564594.

36. Taneera J, Lang S, Sharma A, Fadista J, Zhou Y, Ahlqvist E, et al. A systems genetics approach identifies genes and pathways for type 2 diabetes in human islets. Cell Metab. 2012;16(1):122–34. doi: 10.1016/j.cmet.2012.06.006. PubMed PMID: 22768844.

37. Yu H, Kim PM, Sprecher E, Trifonov V, Gerstein M. The importance of bottlenecks in protein networks: correlation with gene essentiality and expression dynamics. PLoS Comput Biol. 2007;3(4):e59. Epub 2007/04/24. doi: 06-PLCB-RA-0302R2 [pii] 10.1371/journal.pcbi.0030059. PubMed PMID: 17447836; PubMed Central PMCID: PMC1853125.

38. Kadowaki T, Fujita-Yamaguchi Y, Nishida E, Takaku F, Akiyama T, Kathuria S, et al. Phosphorylation of tubulin and microtubule-associated proteins by the purified insulin receptor kinase. J Biol Chem. 1985;260(7):4016–20. Epub 1985/04/10. PubMed PMID: 3920212.

39. Wandosell F, Serrano L, Avila J. Phosphorylation of alpha-tubulin carboxyl-terminal tyrosine prevents its incorporation into microtubules. J Biol Chem. 1987;262(17):8268–73. Epub 1987/06/15. PubMed PMID: 3036806.

40. Sawai M, Uchida Y, Ohno Y, Miyamoto M, Nishioka C, Itohara S, et al. The 3- hydroxyacyl-CoA dehydratases HACD1 and HACD2 exhibit functional redundancy and are active in a wide range of fatty acid elongation pathways. J Biol Chem. 2017;292(37):15538–51. doi: 10.1074/jbc.M117.803171. PubMed PMID: 28784662; PubMed Central PMCID: PMCPMC5602410.

41. Courilleau D, Chastre E, Sabbah M, Redeuilh G, Atfi A, Mester J. B-ind1, a novel mediator of Rac1 signaling cloned from sodium butyrate-treated fibroblasts. J Biol Chem. 2000;275(23):17344–8. Epub 2000/04/05. doi: 10.1074/jbc.M000887200 M000887200 [pii]. PubMed PMID: 10747961.

42. Faure R, Baquiran G, Bergeron JJ, Posner BI. The dephosphorylation of insulin and epidermal growth factor receptors. Role of endosome-associated phosphotyrosine phosphatase(s). J Biol Chem. 1992;267(16):11215–21. PubMed PMID: 1375938.

43. Wandinger-Ness A, Zerial M. Rab proteins and the compartmentalization of the endosomal system. Cold Spring Harb Perspect Biol. 2014;6(11):a022616. Epub 2014/10/25. doi: cshperspect.a022616 [pii] 10.1101/cshperspect.a022616. PubMed PMID: 25341920.

44. Overmeyer JH, Maltese WA. Tyrosine phosphorylation of Rab proteins. Methods Enzymol. 2005;403:194–202. Epub 2006/02/14. doi: S0076-6879(05)03016-8 [pii] 10.1016/S0076-6879(05)03016-8. PubMed PMID: 16473587.

45. Progida C, Malerod L, Stuffers S, Brech A, Bucci C, Stenmark H. RILP is required for the proper morphology and function of late endosomes. J Cell Sci. 2007;120(Pt 21):3729–37. doi: 10.1242/jcs.017301. PubMed PMID: 17959629.

46. De Luca M, Cogli L, Progida C, Nisi V, Pascolutti R, Sigismund S, et al. RILP regulates vacuolar ATPase through interaction with the V1G1 subunit. J Cell Sci. 2014;127(Pt 12):2697–708. doi: 10.1242/jcs. 142604. PubMed PMID: 24762812.

47. Choi E, Zhang X, Xing C, Yu H. Mitotic Checkpoint Regulators Control Insulin Signaling and Metabolic Homeostasis. Cell. 2016. Epub 2016/07/05. doi: S0092- 8674(16)30721-8 [pii] 10.1016/j.cell.2016.05.074. PubMed PMID: 27374329.

48. Balbis A, Baquiran G, Dumas V, Posner BI. Effect of inhibiting vacuolar acidification on insulin signaling in hepatocytes. J Biol Chem. 2004;279(13):12777–85. Epub 2003/12/23. doi: 10.1074/jbc.M311493200 M311493200 [pii]. PubMed PMID: 14688247.

49. Voight BF, Scott LJ, Steinthorsdottir V, Morris AP, Dina C, Welch RP, et al. Twelve type 2 diabetes susceptibility loci identified through large-scale association analysis. Nat Genet. 2010;42(7):579–89. Epub 2010/06/29. doi: ng.609 [pii] 10.1038/ng.609. PubMed PMID: 20581827; PubMed Central PMCID: PMC3080658.

50. Morris AP, Voight BF, Teslovich TM, Ferreira T, Segre AV, Steinthorsdottir V, et al. Large-scale association analysis provides insights into the genetic architecture and pathophysiology of type 2 diabetes. Nat Genet. 2012;44(9):981–90. Epub 2012/08/14. doi: ng.2383 [pii] 10.1038/ng.2383. PubMed PMID: 22885922; PubMed Central PMCID: PMC3442244.

51. Sherr CJ, Roberts JM. Living with or without cyclins and cyclin-dependent kinases. Genes Dev. 2004;18(22):2699–711. PubMed PMID: 15545627.

52. Barriere C, Santamaria D, Cerqueira A, Galan J, Martin A, Ortega S, et al. Mice thrive without Cdk4 and Cdk2. Mol Oncol. 2007;1(1):72–83. Epub 2007/06/01. doi: S1574-7891(07)00011-7 [pii] 10.1016/j.molonc.2007.03.001. PubMed PMID: 19383288.

53. Kim SY, Lee JH, Merrins MJ, Gavrilova O, Bisteau X, Kaldis P, et al. Loss of Cyclin Dependent Kinase 2 in the Pancreas Links Primary beta-cell Dysfunction to Progressive Depletion of beta-cell Mass and Diabetes. J Biol Chem. 2017. doi: 10.1074/jbc.M116.754077. PubMed PMID: 28100774.

54. Rothman JE. The future of Golgi research. Mol Biol Cell. 2010;21(22):3776–80. Epub 2010/11/17. doi: 21/22/3776 [pii] 10.1091/mbc.E10-05-0418. PubMed PMID: 21079007; PubMed Central PMCID: PMC2982129.

55. De Matteis MA, Luini A. Exiting the Golgi complex. Nat Rev Mol Cell Biol. 2008;9(4):273–84. Epub 2008/03/21. doi: nrm2378 [pii] 10.1038/nrm2378. PubMed PMID: 18354421.

56. Gaulin JF, Fiset A, Fortier S, Faure RL. Characterization of Cdk2-cyclin E complexes in plasma membrane and endosomes of liver parenchyma. Insulin-dependent regulation. J Biol Chem. 2000;275(22):16658–65. Epub 2000/05/29. doi: 275/22/16658 [pii]. PubMed PMID: 10828061.

57. Haj FG, Verveer PJ, Squire A, Neel BG, Bastiaens PI. Imaging sites of receptor dephosphorylation by PTP1B on the surface of the endoplasmic reticulum. Science. 2002;295(5560):1708–11. Epub 2002/03/02. doi: 10.1126/science.1067566 295/5560/1708 [pii]. PubMed PMID: 11872838.

58. Grecco HE, Schmick M, Bastiaens PI. Signaling from the living plasma membrane. Cell. 2011;144(6):897–909. doi: 10.1016/j.cell.2011.01.029. PubMed PMID: 21414482.

59. Baumdick M, Bruggemann Y, Schmick M, Xouri G, Sabet O, Davis L, et al. EGFdependent re-routing of vesicular recycling switches spontaneous phosphorylation suppression to EGFR signaling. Elife. 2015;4. Epub 2015/11/27. doi: 10.7554/eLife.12223. PubMed PMID: 26609808; PubMed Central PMCID: PMC4716840.

60. Levy ED, Landry CR, Michnick SW. Cell signaling. Signaling through cooperation. Science. 2010;328(5981):983–4. Epub 2010/05/22. doi: 328/5981/983 [pii] 10.1126/science. 1190993. PubMed PMID: 20489011.

61. Wang W, Fridman A, Blackledge W, Connelly S, Wilson IA, Pilz RB, et al. The phosphatidylinositol 3-kinase/akt cassette regulates purine nucleotide synthesis. J Biol Chem. 2009;284(6):3521–8. Epub 2008/12/11. doi: M806707200 [pii] 10.1074/jbc.M806707200. PubMed PMID: 19068483; PubMed Central PMCID: PMC2635033.

62. Asby DJ, Cuda F, Beyaert M, Houghton FD, Cagampang FR, Tavassoli A. AMPK Activation via Modulation of De Novo Purine Biosynthesis with an Inhibitor of ATIC Homodimerization. Chem Biol. 2015;22(7):838–48. Epub 2015/07/07. doi: S1074- 5521(15)00234-3 [pii] 10.1016/j.chembiol.2015.06.008. PubMed PMID: 26144885.

63. Zhang CS, Jiang B, Li M, Zhu M, Peng Y, Zhang YL, et al. The lysosomal v-ATPase-Ragulator complex is a common activator for AMPK and mTORC1, acting as a switch between catabolism and anabolism. Cell Metab. 2014;20(3):526–40. Epub 2014/07/09. doi: S1550-4131(14)00279-4 [pii] 10.1016/j.cmet.2014.06.014. PubMed PMID: 25002183.

64. Walter P, Ron D. The unfolded protein response: from stress pathway to homeostatic regulation. Science. 2011;334(6059):1081–6. Epub 2011/11/26. doi: 334/6059/1081 [pii] 10.1126/science. 1209038. PubMed PMID: 22116877.

65. Shanik MH, Xu Y, Skrha J, Dankner R, Zick Y, Roth J. Insulin resistance and hyperinsulinemia: is hyperinsulinemia the cart or the horse? Diabetes Care. 2008;31 Suppl 2:S262–8. Epub 2008/02/15. doi: 31/Supplement_2/S262 [pii] 10.2337/dc08-s264. PubMed PMID: 18227495.

66. Ader M, Stefanovski D, Kim SP, Richey JM, Ionut V, Catalano KJ, et al. Hepatic insulin clearance is the primary determinant of insulin sensitivity in the normal dog. Obesity (Silver Spring). 2014;22(5):1238–45. Epub 2013/10/15. doi: 10.1002/oby.20625. PubMed PMID: 24123967; PubMed Central PMCID: PMC3969862.

67. Bergeron JJ, Au CE, Desjardins M, McPherson PS, Nilsson T. Cell biology through proteomics--ad astra per alia porci. Trends Cell Biol. 2010;20(6):337–45. Epub 2010/03/17. doi: S0962-8924(10)00038-3 [pii] 10.1016/j.tcb.2010.02.005. PubMed PMID: 20227883.

68. Ferrannini E. The target of metformin in type 2 diabetes. N Engl J Med. 2014;371(16):1547–8. Epub 2014/10/16. doi: 10.1056/NEJMcibr1409796. PubMed PMID: 25317875.

69. Havlis J, Thomas H, Sebela M, Shevchenko A. Fast-response proteomics by accelerated in-gel digestion of proteins. Anal Chem. 2003;75(6):1300–6. PubMed PMID: 12659189.

70. Shevchenko A, Wilm M, Vorm O, Mann M. Mass spectrometric sequencing of proteins silver-stained polyacrylamide gels. Anal Chem. 1996;68(5):850–8. PubMed PMID: 8779443.

71. Au CE, Bell AW, Gilchrist A, Hiding J, Nilsson T, Bergeron JJ. Organellar proteomics to create the cell map. Curr Opin Cell Biol. 2007;19(4):376–85. Epub 2007/08/11. doi: S0955-0674(07)00096-8 [pii] 10.1016/j.ceb.2007.05.004. PubMed PMID: 17689063.

72. Yu H, Braun P, Yildirim MA, Lemmens I, Venkatesan K, Sahalie J, et al. High-quality binary protein interaction map of the yeast interactome network. Science. 2008;322(5898):104–10. Epub 2008/08/23. doi: 1158684 [pii] 10.1126/science.1158684. PubMed PMID: 18719252; PubMed Central PMCID: PMC2746753.

73. Mellacheruvu D, Wright Z, Couzens AL, Lambert JP, St-Denis NA, Li T, et al. The CRAPome: a contaminant repository for affinity purification-mass spectrometry data. Nat Methods. 2013;10(8):730–6. doi: 10.1038/nmeth.2557. PubMed PMID: 23921808; PubMed Central PMCID: PMCPMC3773500.

74. Morone F, Makse HA. Influence maximization in complex networks through optimal percolation. Nature. 2015;524(7563):65–8. Epub 2015/07/02. doi: nature14604 [pii] 10.1038/nature 14604. PubMed PMID: 26131931.

75. Xue Y, Ren J, Gao X, Jin C, Wen L, Yao X. GPS 2.0, a tool to predict kinase-specific phosphorylation sites in hierarchy. Mol Cell Proteomics. 2008;7(9):1598–608. doi: 10.1074/mcp.M700574-MCP200. PubMed PMID: 18463090; PubMed Central PMCID: PMCPMC2528073.

76. Hornbeck PV, Zhang B, Murray B, Kornhauser JM, Latham V, Skrzypek E. PhosphoSitePlus, 2014: mutations, PTMs and recalibrations. Nucleic Acids Res. 2015;43(Database issue):D512–20. doi: 10.1093/nar/gku1267. PubMed PMID: 25514926; PubMed Central PMCID: PMCPMC4383998.

77. Linding R, Jensen LJ, Pasculescu A, Olhovsky M, Colwill K, Bork P, et al. NetworKIN: a resource for exploring cellular phosphorylation networks. Nucleic Acids Res. 2008;36(Database issue):D695–9. doi: 10.1093/nar/gkm902. PubMed PMID: 17981841; PubMed Central PMCID: PMCPMC2238868.

78. Boyle AP, Hong EL, Hariharan M, Cheng Y, Schaub MA, Kasowski M, et al. Annotation of functional variation in personal genomes using RegulomeDB. Genome Res. 2012;22(9):1790–7. doi: 10.1101/gr.137323.112. PubMed PMID: 22955989; PubMed Central PMCID: PMCPMC3431494.

79. van Dam S, Craig T, de Magalhaes JP. GeneFriends: a human RNA-seq-based gene and transcript co-expression database. Nucleic Acids Res. 2015;43(Database issue):D1124–32. doi: 10.1093/nar/gku1042. PubMed PMID: 25361971; PubMed Central PMCID: PMCPMC4383890.

80. Mootha VK, Lindgren CM, Eriksson KF, Subramanian A, Sihag S, Lehar J, et al. PGC-1alpha-responsive genes involved in oxidative phosphorylation are coordinately downregulated in human diabetes. Nat Genet. 2003;34(3):267–73. doi: 10.1038/ng1180. PubMed PMID: 12808457.

81. Subramanian A, Tamayo P, Mootha VK, Mukherjee S, Ebert BL, Gillette MA, et al. Gene set enrichment analysis: a knowledge-based approach for interpreting genome-wide expression profiles. Proc Natl Acad Sci U S A. 2005;102(43):15545–50. doi: 10.1073/pnas.0506580102. PubMed PMID: 16199517; PubMed Central PMCID: PMCPMC1239896.

